# Hippocampal subfields and limbic white matter jointly predict learning rate in older adults

**DOI:** 10.1101/661702

**Authors:** Andrew R. Bender, Andreas M. Brandmaier, Sandra Düzel, Attila Keresztes, Ofer Pasternak, Ulman Lindenberger, Simone Kühn

**Affiliations:** Departments of Epidemiology and Biostatistics, Neurology and Ophthalmology, Michigan State University, East Lansing, MI, USA; Center for Lifespan Psychology, Max Planck Institute for Human Development, & Max Planck UCL Centre for Computational Psychiatry and Ageing Research, Berlin, Germany and London, UK; Center for Lifespan Psychology, Max Planck Institute for Human Development, Berlin, Germany; Center for Lifespan Psychology, Max Planck Institute for Human Development, Berlin, Germany Research Centre for Natural Sciences, Hungarian Academy of Sciences, Budapest, Hungary, Faculty of Education and Psychology, Eotvos Lorand University, Budapest, Hungary; Departments of Psychiatry and Radiology, Brigham and Women’s Hospital, Harvard Medical School, Boston, MA, USA; Center for Lifespan Psychology, Max Planck Institute for Human Development, & Max Planck UCL Centre for Computational Psychiatry and Ageing Research, Berlin, Germany, and European University Institute, San Domenico di Fiesole (FI), Italy; Center for Lifespan Psychology, Max Planck Institute for Human Development, & Department of Psychiatry and Psychotherapy, University Clinic Hamburg-Eppendorf, Hamburg, Germany

**Keywords:** hippocampus, white matter, aging, verbal learning, memory, Berlin Aging Study II

## Abstract

Age-related memory impairments have been linked to differences in structural brain parameters, including cerebral white matter (WM) microstructure and hippocampal (HC) volume, but their combined influences are rarely investigated. In a population-based sample of 337 older participants 61–82 years of age (M_age_=69.66, SD_age_=3.92 years) we modeled the independent and joint effects of limbic WM microstructure and HC subfield volumes on verbal learning. Participants completed a verbal learning task over five learning trials and underwent magnetic resonance imaging (MRI), including structural and diffusion scans. We segmented three HC subregions on high-resolution MRI data and sampled mean fractional anisotropy (FA) from bilateral limbic WM tracts identified via deterministic fiber tractography. Using structural equation modeling, we evaluated the associations between learning rate and latent factors representing FA sampled from limbic WM tracts, and HC subfield volumes, as well as their latent interaction. Results showed limbic WM and the interaction of HC and WM – but not HC volume alone – predicted verbal learning rates. Model decomposition revealed HC volume is only positively associated with learning rate in individuals with higher levels of WM anisotropy. We conclude that structural characteristics of limbic WM regions and HC volume jointly contribute to verbal learning in older adults.

## Hippocampal Subfields and Limbic White Matter Jointly Predict Learning Rate in Older Adults

Age-related deficits in verbal learning and memory have a long history of study in psychological and cognitive sciences (Kausler, 1994; Korchin and Basowitz, 1957; Salthouse, 1985). Individual differences in learning and memory in older adults are linked with differences in both regional gray matter volumes in the medial temporal lobe (MTL; Petersen et al., 2000) and microstructural measures of limbic white matter (WM) pathways (see Madden et al., 2012 for a review). However, few studies to date have investigated the influences of both classes of neuroanatomical correlates of verbal learning, using two magnetic resonance imaging (MRI) modalities, in a population-based cohort of older adults. Arguably, modeling learning and memory as a function of a larger, integrated neural system affords a more balanced perspective over traditional univariate modeling of individual neural structures (Aggleton, 2014).

Verbal learning is commonly tested via serial presentation of lexical stimuli, followed by tests of free or cued recall and recognition. For instance, multiple neuropsychological instruments assessing verbal learning repeatedly present the same stimulus list over multiple successive learning trials; following each presentation, participants freely recall as many items as possible (Baldo et al., 2002; Schmidt, 1996). Although the sum of recalled items is commonly used as measure of aggregate performance, the slope of change in memory performance across learning trials can serve as an estimate of the rate of learning (Jones et al., 2005). Although many studies have explored the neural correlates of age-associated decrements in delayed mnemonic retrieval, fewer have investigated the structural brain correlates associated with learning rate (Gifford et al., 2015). These initial studies suggest that individual differences in learning *rate* are associated with hippocampal (HC) volumes in normal aging and mild cognitive impairment (Bonner-Jackson et al., 2015; Gifford et al., 2015), and may afford a more sensitive behavioral correlate of brain organization. Although prior work has linked larger hippocampal subregional volumes in Cornu Ammonis (CA) and dentate gyrus (DG) to better verbal learning, associative memory, and higher longitudinal retest improvements (Bender et al., 2013; Mueller et al., 2011; Shing et al., 2011), this has not been investigated as a correlate of learning rate.

In addition to HC, other established neuroanatomical correlates of age-associated memory decline include structurally linked afferent and efferent limbic WM pathways measured via diffusion magnetic resonance imaging (dMRI; Bender et al., 2016; Bennett et al., 2014; Charlton et al., 2013; Fletcher et al., 2013; Henson et al., 2016; Metzler-Baddeley et al., 2011a; Sasson et al., 2013; Sepulcre et al., 2008; Stoub et al., 2006; see Preston and Eichenbaum, 2013; Shing et al., 2008 for reviews). In particular, these extant reports implicate cingulum bundle, including dorsal and parahippocampal segments, fornix, and uncinate fasciculus (UF) as primary WM correlates of age-related memory declines. Furthermore, associations between total HC volume and memory in older adults are inconsistent (Van Petten, 2004), and suggest other, less elucidated factors may modify this relationship. One possibility is that its structure includes multiple functionally and cytoarchitectonically distinct subregions, which show differential associations with aging and with mnemonic processes (Braak et al., 1996; Duvernoy, 2005; Insausti et al., 1998; Kiernan, 2012; Wilson et al., 2006). This perspective suggests that the relationship between total HC volume and memory may be attenuated by the lack of functional and structural specificity (Van Petten, 2004). Alternatively, associations between HC volume and memory may be modified by related factors, such as the extent of WM connectivity (Foster et al., 2019; Metzler-Baddeley et al., 2019). However, the statistical interaction between HC volumes and limbic WM microstructure has not been tested previously.

Investigating the links of different HC subfields and limbic WM fiber tracts to episodic memory requires a sufficiently large sample and the incorporation of multiple MRI measurement approaches. The Berlin Aging Study-II (BASE-II; Bertram et al., 2014; Gerstorf et al., 2016) includes over 1500 healthy older adults; of these, a subset of whom underwent MRI neuroimaging. The sample is sufficiently large to permit the use of structural equation modeling (SEM), which allows the confirmatory construction of memory performance and brain parameters as latent factors, moving from an observed level to a more valid construct level. These latent factors then serve as a basis for exploring brain–behavior associations as between– construct correlations, independent of measurement error. Thus, following the plea of Brandmaier et al. (2013), we use SEM as a statistical tool that combines the benefits of both confirmatory and explanatory modes of scientific inquiry.

Specifically, we were interested in modeling the rate of learning across five free recall trials of a test of verbal learning (Helmstaedter and Durwen, 1989; Schmidt, 1996) to test the notion that rate of learning is a more sensitive correlate of brain structure and organization than the intercept. By evaluating the combined contributions from both limbic WM and HC subfield volumes under the SEM framework, our goal was to simultaneously model multiple distinct, but interdependent structural neural correlates of learning. Our initial approach was more exploratory by freely estimating all associations between individual HC and WM factors and verbal learning slope and intercept. We expected a similar pattern of anatomical associations with learning as previously reported: higher FA in cingulum and fornix tracts and larger hippocampal volumes, particularly of CA and DG (Bender et al., 2016; Bennett et al., 2014; Madden et al., 2012). Herein, we also tested the general hypothesis that aggregate, latent measures of HC volume and limbic white matter fractional anisotropy in older adults are both associated with verbal learning rate. Furthermore, we hypothesized that individual differences in rates of learning would be associated with individual differences in the interaction between HC volume and FA in limbic WM.

## Methods

### Participants

Study data were drawn from the first wave of the BASE-II cohort (Bertram et al., 2014; Gerstorf et al., 2016), a population-based study of older and younger adults living in Berlin, Germany. The baseline cohort included 1532 older adults from 60 to 88 years of age. None of the participants took medication that might affect memory function, and none had neurological disorders, psychiatric disorders, or a history of head injuries. All participants reported normal or corrected to normal vision and were right-handed. All participants were invited to two cognitive sessions with an exact interval of seven days and at the same time of day to avoid circadian confounding effects on session differences in performance. Participants were tested in groups of 4–6 individuals. The ethics section of the German Psychological Society approved the study (SK 012013_6). All participants had provided informed consent in accord with the Declaration of Helsinki.

After completing the comprehensive cognitive assessment in BASE-II, MR-eligible participants were invited to take part in one MRI session within a mean time interval of 3.2 months after the cognitive testing. The subsample consisted of 345 older adults aged 61–82 years (mean age 70.1 years, SD = 3.9 years, 39% female). We excluded six participants following technical errors in cognitive test administration, and we excluded two additional participants with scores below 25 on the Mini-Mental Status Examination (MMSE; Folstein et al., 1975). Most participants’ MMSE scores were well above this cut-off (mean = 28.61, SD = 1.15). BASE-II participants in the MRI cohort did not differ from those who did not undergo MRI scanning in terms of educational attainment, cognitive performance, or MMSE scores (for all, *t* < 1.0), although the MRI cohort was significantly younger than the non-MRI cohort (*t* = 2.577, *p* < .05) by approximately 6 months. The final sample retained for analysis included 337 older adults (mean age = 69.66, SD = 3.92 years). Sample demographics showed a greater proportion of men (61.7%) than women (38.3%), and mean level of years of education nearing one year of university (mean = 14.07, SD = 2.90 years).

### Magnetic Resonance Imaging (MRI)

#### Image acquisition

All MRI data were acquired on a 3T Siemens Magnetom Tim Trio scanner. For most cases, a 32-channel head coil was used, although in two cases a 12-channel coil was used as the 32-channel coil provided an uncomfortable fit. MRI data acquisition included a T1-weighted magnetization-prepared rapid gradient echo (MPRAGE) sequence, acquired in the sagittal plane with a single average, repetition time (TR) = 2,500 ms, echo time (TE) = 4.77 ms, with an isotropic voxel size of 1.0 × 1.0 × 1.0 mm, using a 3D distortion correction filter and pre-scan normalization with FOV = 256, and GRAPPA acceleration factor = 2. Acquisition also included a single T_2_-weighted, turbo spin echo (TSE) high-resolution sequence in a coronal direction, oriented perpendicularly to the long axis of the left HC, with voxel size = 0.4 × 0.4 × 2.0 mm, 30 slices. TR = 8,150 ms, TE = 50 ms, flip angle = 120**°**, positioned to cover the entire extent of the HC. A single-shot, echo-planar imaging, diffusion weighted sequence was also acquired in transverse plane with TR = 11,000 ms, TE = 98 ms, in 60 gradient directions, diffusion weighting of b = 1,000 s/mm^2^, seven volumes collected without diffusion weighting (b = 0) and generalized autocalibrating partially parallel acquisitions (GRAPPA) acceleration factor = 2 with an isotropic voxel of 1.70 mm^3^.

#### Diffusion MRI Processing

All diffusion-weighted images (DWI) underwent an initial quality control (QC) process using DTIPrep v. 1.2.4 (Oguz et al., 2014), software to eliminate noisy gradient directions and correct for motion and eddy currents.

#### Free water estimation and removal

The influence of partial volume artifacts from cerebral spinal fluid (CSF) is an established limitation of the single-tensor diffusion tensor imaging (DTI) model that is often used in the context of tractography (Concha et al., 2005; Metzler-Baddeley et al., 2011b; Pasternak et al., 2009), particularly in regions directly adjacent to ventricular CSF, such as the fornix (Jones and Cercignani, 2010; Metzler-Baddeley et al., 2011a). To address this limitation, tensor data were corrected for CSF contamination on a voxel-by-voxel basis, using the free water elimination MATLAB code (FWE; Pasternak et al., 2009), resulting with free-water corrected diffusion tensors. The free water corrected tensors were then decomposed using FSL to produce FA image maps.

#### Sample template creation

We used DTI-TK (Zhang et al., 2006) software to align all participants’ data into a common template. The complete procedures for inter-subject registration are detailed in Supplementary Materials.

#### Region of Interest (ROI) creation

Individual ROIs reflecting seed regions, regions of inclusion, and regions of exclusion were drawn on the template-space FA image, colored by orientation image output as an option by DTI-TK in ITK-SNAP (www.itksnap.org; Yushkevich et al., 2006). All template-space ROIs were nonlinearly deprojected to native space, where they were inspected for errors by one of the authors (A.R.B.). Following published deterministic tractography approaches for these regions (Bennett et al., 2014; Malykhin et al., 2008; Metzler-Baddeley et al., 2011a), we created ROIs for tractography of four, bilateral limbic WM: dorsal cingulum bundle (CBD), parahippocampal cingulum bundle (CBH), posterior fornix, and uncinate fasciculus (UF). The procedures for tractography mask creation, placement, and spatial transformation from standard to native space are detailed in Supplementary Materials.

#### Constrained spherical deconvolution (CSD) tractography

To enhance the anatomical validity and minimize the potentially confounding influences of crossing fiber populations, we performed diffusion MR tractography using constrained spherical deconvolution (CSD), a method that fits a fiber orientation distribution (FOD) to each voxel and performs tractography based on peaks in the FOD (Tournier et al., 2007; Tournier et al., 2004). CSD tractography is considered superior to other commonly used approaches for delineating WM tracts of interest, such as white matter skeletonization, due to greater anatomical precision for any given tract (Metzler-Baddeley et al., 2011a). MRtrix3 (Tournier et al., 2007; Tournier et al., 2004) software was used for CSD-based deterministic tractography on the DWI data following QC, but before any free water correction. Response function estimation used the method previously described (Tournier et al., 2013) with maximum spherical harmonic degree = 4. Following default procedures (“Beginner DWI Tutorial,” 2017), two separate FOD images were produced using the estimated response function, one using a whole brain mask, and the other using the thresholded FA mask (here FA prior to free water elimination was used). The whole brain mask was used only for tractography of the fornix, as using the thresholded FA masked FOD data did not permit sufficient information for reliable tractography of fornix, whereas use of the whole brain mask for other regions produced excessive spurious and anatomically implausible streamlines (i.e., crossing sulci). For fornix, a secondary mask was also applied, in which the operator (A.R.B.) drew an inclusionary ROI to cover the fornix, but exclude other regions of the ventricles, the thalamus, and choroid plexus. Additional exclusionary masks were liberally applied outside these regions to limit any spurious streamlines. The thresholded, FA-masked FOD data were used for CSD tractography of the other regions: CBD CBH, and UF. All streamline outputs were inspected by the same person (A.R.B.) using the MRtrix image viewer to ensure complete inclusion and to check for spurious streamlines (see Fig. SM1 for an example of sampled streamlines). Additional information on streamline inspection is available in Supplementary Materials. Following streamline generation and inspection, we used streamlines to sample median free-water corrected FA values, and then calculated the mean value across all streamlines in each tract of interest.

#### Hippocampal subfield morphometry

HC subfield regions included were based on methods from prior work (Bender et al., 2013; Daugherty et al., 2016; Mueller et al., 2011; Mueller et al., 2007; Shing et al., 2011), and included separate regions for SUB, and aggregations of CA1 and 2 (CA1/2), and an aggregation of CA3 and the DG (CA3/DG).

#### Optimized automated segmentation

We used the Automated Segmentations of Hippocampal Subfields (ASHS; Yushkevich et al., 2015a; Yushkevich et al., 2010) software with a customized atlas for HC subfield morphometry (Bender et al., 2018). Because it uses multi-atlas label fusion methods, ASHS may be more sensitive to individual differences in HC subfield morphology, than single-atlas approaches, such as Freesurfer. The customized atlas was built using a modified version of the manual demarcation and tracing rules described previously (Daugherty et al., 2016; Keresztes et al., 2017; see Supplementary Materials for more information).

#### ICV correction

We sampled the intra-cranial vault (ICV) as described previously (Bender et al., 2013; Keihaninejad et al., 2010), using the brain extraction tool (BET; Smith, 2002) in FSL 5.0 (Jenkinson et al., 2005; 2012) on the MPRAGE images (further detail is provided in Supplementary Materials).

### Cognitive Testing

The Verbal Learning and Memory Test (VLMT; Helmstaedter and Durwen, 1989) is a German version of the Rey Auditory Verbal Learning Test (Schmidt, 1996). Participants heard a list of 15 nouns, serially presented via headphones. Presentation of the list was followed by a recall phase in which a computer screen prompted the participants to type as many words as they could remember from the list. This was repeated over five learning trials, each with the same word list. The same list of German nouns was used for all participants. The present analyses are based on the five verbal learning trials and do not include data from the delayed recall and recognition tasks that are also part of the VLMT.

### Data Conditioning

ICV values were divided by 1,000 to align the scales of HC subfields and ICV, and to increase numerical stability in the parameter estimation. We corrected each of the subregional HC volumes for ICV using the analysis of covariance approach (Bender et al., 2013; Jack et al., 1989; Raz et al., 2005). All analyses reported below used the adjusted HC subfield volumes. In addition, all FA values were centered at their respective sample means and HC subfield volumes were standardized to z-scores.

### Data Analysis

#### Overview

Data modeling and analysis was performed in Mplus 7 (Muthén and Muthén, 2012). We used latent factor analysis to explore associations among verbal learning, HC subfield volumes, and WM FA within an overall multivariate model. In light of the complex Model specification proceeded in an iterative fashion, as we sought to establish the validity of each. The first step involved specifying, testing, and refining individual measurement models for latent variables (factors) in each domain, including: 1) a latent growth model across learning trials (specified below) yielded factors for intercept and slope, 2) three separate HC subfield volume factors, and 3) four latent factors representing mean FA of four WM tracts of interest (see Table SM1 for factor loadings). Following this, we combined these individual measurement models with a structural model to test associations across domains between all latent factors (Fig. 1).

**Figure 1.**
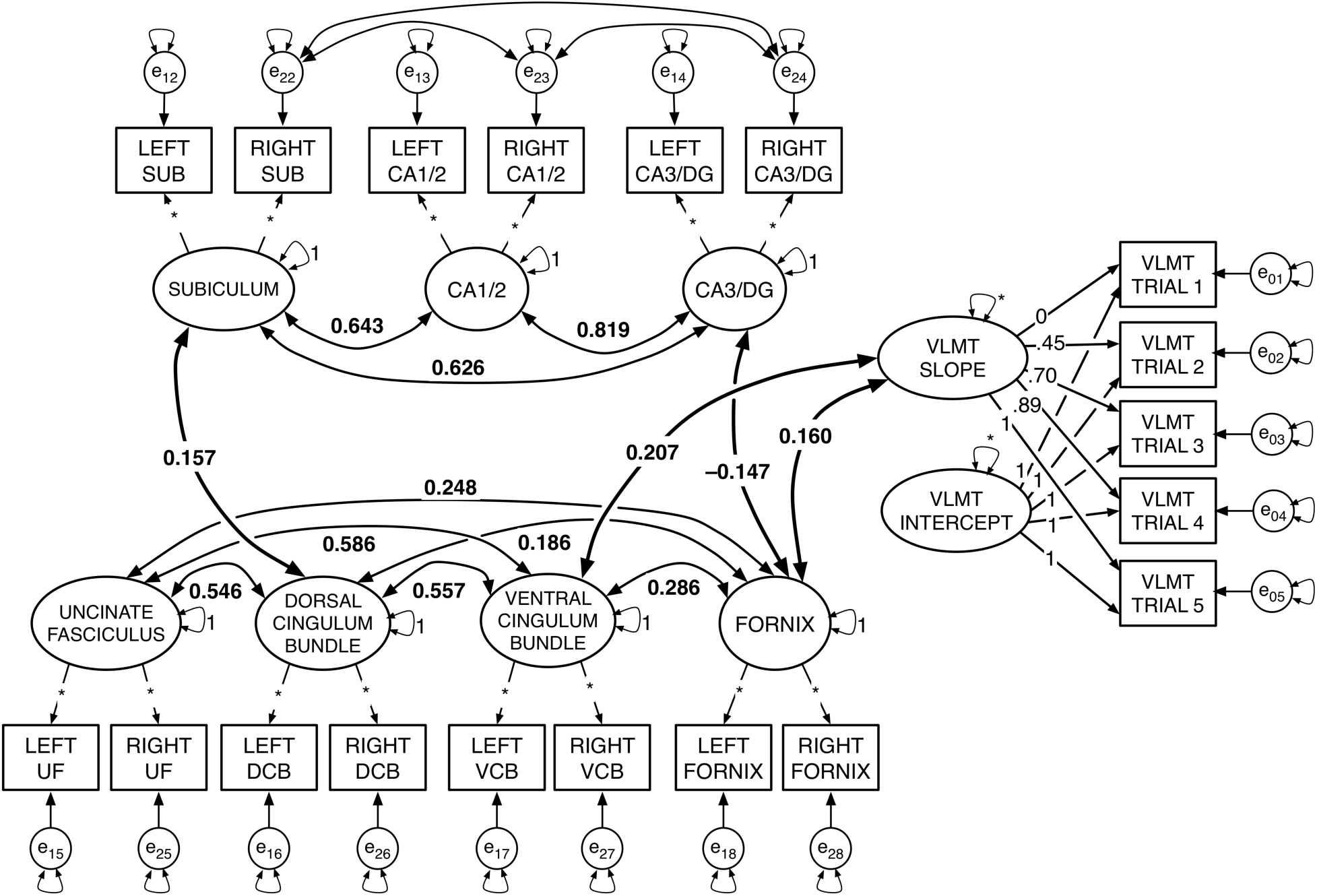
Diagram of the ‘combined model,’ with only significant correlations (i.e., *p* < .05) shown. The estimated model included the fully saturated latent correlation matrix. Larger ellipses with small double-headed arrows represent latent factors with variance either fixed at 1 or freely estimated (*). Small single-headed arrows between factors and their respective observed indicators reflect factor loadings, with the value of the factor loading pre-specified (i.e., based on estimated loadings from earlier modeling steps) or freely estimated (*). The rectangles reflect the observed indicators for each measurement, and the small circles with double-headed arrows reflect their residuals and residual variance; all residual error variance parameters were freely estimated. In the LGM portion on the right side of the figure, (right) factor loadings for the LGM slope factor were originally estimated using a latent basis free model. The intercept (i.e., all factor loadings fixed to 1), and slope with factor loadings represent individual differences in growth across the verbal learning trials. (See Table SM3 for a comparison for factor loadings and fit to a model with linear slope). Larger curved bidirectional arrows represent significant covariances between factors (*p* < .05), and covariance path values reflect standardized parameters. The figure shows only the significant associations within each brain domain (i.e., among HC subfield factors, and among factors for WM tract FA), and between WM and HC factors. In addition, the bold covariance double-headed arrows show significant associations between brain factors and the slope factor for the LGM on learning trials.

Next, we specified an additional model to test the fit of separate second-order factors representing the HC and WM factors (Fig. 2A). Second-order factors are constructed from other estimated factors, rather than observed indicators. Thus, this approach permits estimation of overall HC and WM factors based on their constituent factors by subregion or tract, which should provide more reliable brain factor estimates. We then tested regression paths from both second-order factors for HC and WM to the learning slope factor. Last, we modeled the latent interaction (Fürst & Ghisletta, 2009; Maslowsky, Jager, & Hemken, 2014; Little, Bovaird, & Widaman, 2006) between the second-order factors for HC and WM to test the hypothesis that the association between HC volume and learning is modified by WM microstructure.

**Figure 2.**
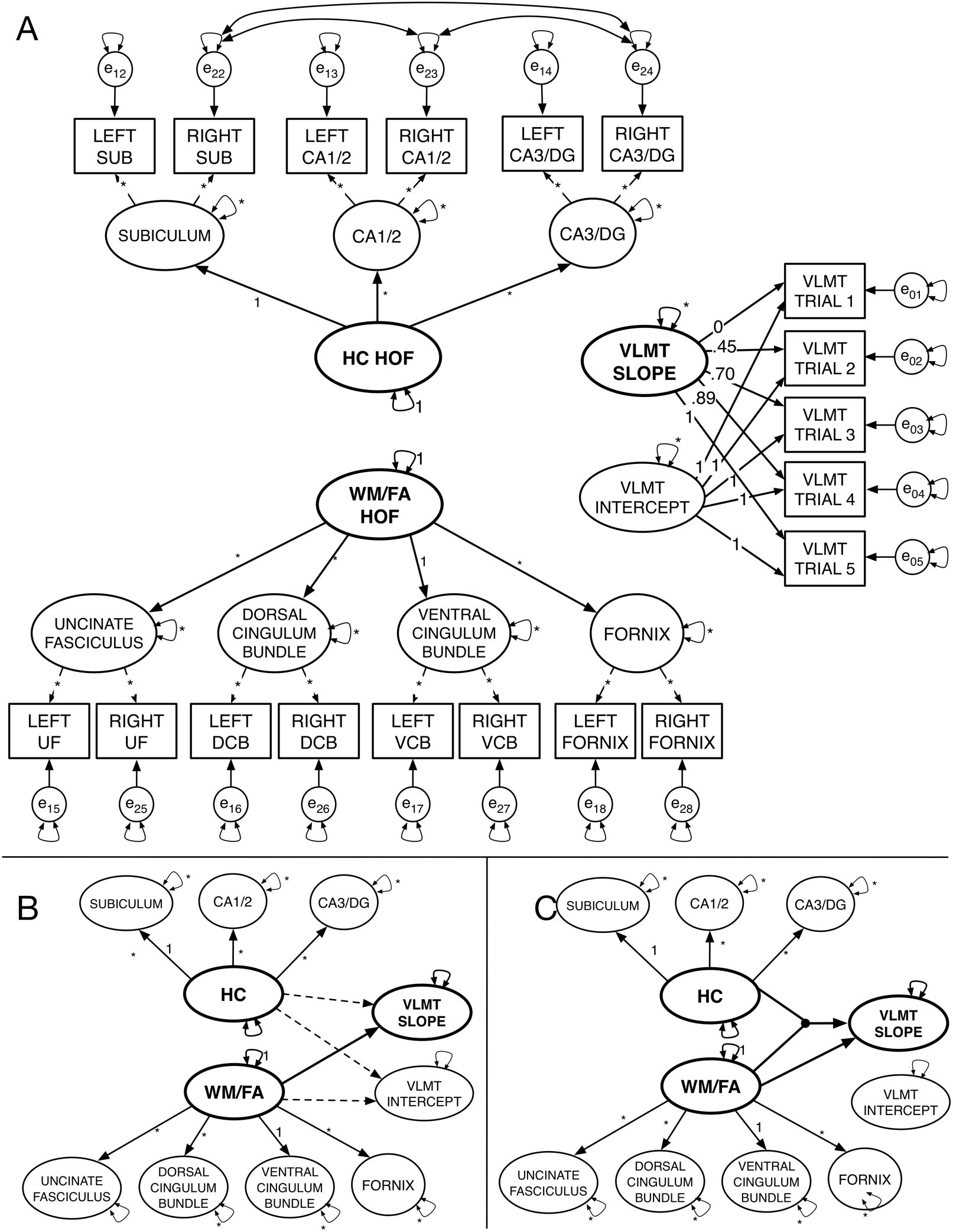
Data modeling steps for second-order models and latent moderated structural equation models. A. Initial specification of the second-order latent factor model. No covariances or regression paths between factors are illustrated. B. Reduced illustration of the specified model (indicators and error variances not shown). Initial specification without latent interaction included regression paths from the HC and WM/FA second-order factors to the slope and intercept factors. Dashed lines indicate nonsignificant regression paths, and solid lines reflect significant paths. C. The latent interaction model showing significant paths from HC and WM/FA factors to the slope factor. The dot symbolizes the latent interaction between the HC and WM/FA factors.

All models used full information maximum likelihood (FIML) to account for missing data without requiring pairwise deletion. Goodness of model fit was assessed by multiple indices including chi-square (χ^2^), chi-square value divided by degrees of freedom (χ^2^/*df*), comparative fit index (CFI), root mean square error of approximation (RMSEA), and standardized root mean square residual (SRMR). We evaluated models according to commonly accepted goodness-of-fit thresholds, that is, non-significant chi-square values, CFI values > 0.95, RMSEA values reliably < 0.05, and SRMR values < 0.05 indicate good model fit (Bentler, 1990; Hooper et al., 2008).

#### Verbal learning latent growth model

We modeled verbal learning by fitting a latent free basis model (McArdle, 1986), a type of latent growth model (LGM; Duncan et al., 2013; Meredith and Tisak, 1990) to the five learning trials. LGM is commonly used to assess latent change over successive occasions, and generally includes separate factors for the intercept and for change trajectories. LGM commonly specifies fixed factor loadings in incrementing value following a specific change function (i.e., linear, quadratic, etc.) across sequential indicators. In contrast, the latent free basis model fixes the factor loadings of the first and last trials at one and zero, respectively; the other factor loadings are freely estimated. Descriptions of the full modeling procedures for the LGM are reported in Supplementary Materials. Briefly, however, it should be noted that initial attempts to fit the LGM with only intercept and linear slopes were poor fit for the data (Table SM3). However, latent basis models may be the best approaches for estimating relative change over trials when linear, quadratic, cubic or other functional patterns do not provide a good fit for the data (Berlin et al., 2013).

#### WM tract and HC subregional factor models

To model each brain imaging parameter, we used a confirmatory factor analysis (CFA) approach (see Supplementary Materials for a complete description). That is, we fit individual, single-factor models for each of the three HC subfield volumes and for each combination of the four WM tracts. For each single-factor model, left and right hemisphere brain parameter measures (i.e., FA or mm^3^ brain volume) served as dual indicators (Bender and Raz, 2015; Raz et al., 2005). In all dual-indicator models of brain imaging parameters, the factor’s variance parameter was fixed to 1 and the factor loadings for both indicators were freely estimated, as were the residual variances. This approach of separate factors per region was preferred over combining all indicator loading into a single factor (see Supplemental Materials for a complete description).

#### Combined model

Next, we specified a combined model that included: (a) the verbal learning intercept and slope factors; (b) the three factors of HC subfields volume; and (c) four factors of FA in WM tracts. Then we estimated a fully crossed latent covariance matrix for each combined model. We used bootstrapped resampling with 1,000 draws to generate confidence intervals around the combined model parameters, and to test significance of the combined model parameters.

#### Second-order factor model

In the following steps, we re-specified the model to estimate two second-order factors: one representing all four WM factors and one representing the four HC subfield factors (Fig. 2A). This model was initially estimated with covariances freely estimated among factors. Following the observation of good fit, we then respecified the model to include directional regression paths from the factors representing both WM and HC to the LGM slope and intercept factors. Although such regression procedures imply causal relationships, it is worth noting that the data were cross-sectional and thus cannot inform regarding order or directionality of age-related changes (Lindenberger et al., 2011).

#### Latent moderation models

Following successful convergence and estimation of the second-order factor model, we followed published suggestions for modeling the latent interaction (Fürst & Ghisletta, 2009; Maslowsky, Jager, & Hemken, 2014; Little, Bovaird, & Widaman, 2006) *between the factors* for the HC and WM second-order factors. We specified the regression paths from the two latent factors representing the brain (i.e., WM and HC), and from their latent interaction to verbal learning to test if the effect of HC on learning varies across levels of WM, and vice versa. We then compared model fit between the models with and without the latent interaction using log-likelihood ratio tests. In addition, we estimated difference in *R*^2^ and variance accounted for in learning rate with and without estimating the interaction between WM and HC. Next, we applied the Johnson-Neyman (1936; Preacher et al., 2006) technique for plotting the effects of each factor in the interaction, WM and HC, on the learning rate factor, at different levels of the other. That is, we plotted the effects of WM on learning rate at different levels of HC volume, and vice versa. Last, we tested simple slopes of each WM or HC predictor on learning slope for each level of the other.

#### Covariate models

To determine if our findings were influenced by relevant demographic variables, we re-evaluated the combined, second-order factor, and latent interaction models with the inclusion of the covariates age in years, sex, and number of years of formal education. Years of age and education were centered at their respective sample means.

## Results

### Associations between WM, HC, and learning parameters

Initial models for WM and HC showed that individual factor models by subregions or WM tracts fit better than single factors models (Table SM2). Similarly, the latent basis free LGM fit better than modeling learning as a linear slope. Furthermore, combining the verbal learning LGM with the individual factors for the seven factors representing HC subfield volumes and limbic WM tracts also resulted in excellent fit. This combined model estimated associations between the seven factors representing the structural brain parameters – free-water corrected FA in four limbic WM fiber tracts and ICV-corrected volumes in three HC subregions – and verbal learning (Fig. 1).

In the combined model, we found that higher learning rate was significantly associated with higher FA in CBH (std. est. = 0.207, *p* = .002) and fornix (std. est. = 0.160, *p* =.025). No additional significant associations were observed between HC subfield factors and verbal learning. Of note, the intercept factor was not significantly associated with any brain factor or with the slope factor.

### Second-order factor model

Based on the combined model, we also specified a model in which the four WM factors UF, CBD, CBH, and fornix, load onto a second-order factor representing WM, and the three HC subfield factors SUB, CA1/2, and CA3/DG load onto the HC volume second-order factor (Fig. 2A). Following the observation of a nonsignificant relationship between brain factors and the intercept of the LGM, we re-specified the model to only estimate the direct paths between the HC and WM second-order factors and the learning slope factor; this model specification proved a good fit for the data (Table SM2). In addition, the *R*^2^ values output by Mplus showed the second order factors accounted for a large and significant proportion of the variance in their constituent factors (Table SM4). However, the only significant covariance between factors was a positive association between the WM factor representing combined FA and the learning slope parameter (r = .195, *p* = .007). Notably, HC was associated with neither the WM factor nor the learning curve factor. In addition, the LGM intercept factor was also unrelated to other latent factors.

### Latent moderation model

Next, using the latent moderated SEM (LMS) approach as implemented in Mplus, we estimated 1) the latent interaction *between* the HC and WM second-order factors, and 2) the direct regression paths from the HC and WM second-order factors and the latent interaction parameter to the intercept and slope on the LGM (Fürst & Ghisletta, 2009; Maslowsky, Jager, & Hemken, 2014; Little, Bovaird, & Widaman, 2006). The use of the maximum likelihood estimation with robust standard errors (MLR) estimator necessary for LMS in Mplus does not provide standard fit indices for model comparison. Thus, we used log-likelihood ratio tests to compare fit between the models with and without the latent interaction. Modeling the latent moderation effect resulted in a significantly better fit (Table SM6). In addition, we estimated difference in *R*^2^ and variance accounted for in learning rate with and without estimating the interaction between WM and HC and calculated the differences in explained variance using the formula provided by Maslowsky et al. (2015; Table 1). The model with the latent interaction explained an additional 3.6% of variance in verbal learning over models without estimating parameter.

**Table 1.** Differences in explained variance with and without latent interaction

| Model | Original Model | | | Latent Moderation Model | | | | Difference<br>$\Delta R^2$ |
| --- | --- | --- | --- | --- | --- | --- | --- | --- |
| | $\beta_{YX1}$ | $\beta_{YX2}$ | $R^2$ | $\beta_{YX1}$ | $\beta_{YX2}$ | $\beta_{X1X2}$ | $R^2$ | |
| No Covariates | 0.452 | 0.290 | 0.074 | 0.372 | 0.293 | 0.486 | 0.110 | 0.036 |
| Age | 0.426 | 0.030 | 0.038 | 0.358 | 0.048 | 0.479 | 0.077 | 0.040 |
| Age, Sex | 0.416 | 0.019 | 0.036 | 0.297 | -0.050 | 0.461 | 0.060 | 0.025 |
| Age, Sex, Edu | 0.391 | 0.024 | 0.032 | 0.227 | -0.273 | 0.483 | 0.050 | 0.017 |

Note. $R^2$ and $\Delta R^2$ (i.e., change in $R^2$ ) was calculated using the formula provided by Maslowsky et al., (2015). $\beta_{YX1}$ : SEM model parameter for regression path to verbal learning slope latent factor from latent factor WM (X1). $\beta_{YX2}$ : SEM model parameter for regression path to verbal learning slope factor from latent factor HC (X2). $\beta_{X1X2}$ : Covariance between the factors for HC and WM. $R^2$ values reflect only variance in verbal learning slope factor explained by the two latent factors WM (X1) and HC (X2), and by their latent interaction. $\Delta R^2$ : difference in $R^2$ values with and without latent interaction. In covariate models, $R^2$ estimates reflect inclusion of age, sex and education.

Next, we followed the Johnson-Neyman (1936) technique for plotting the effects of each factor in the interaction, WM and HC, on the learning rate factor, at different levels of the other. That is, we plotted the effects of WM on learning rate at different levels of HC volume, and vice versa. Last, we extended this approach to test simple slopes of each WM or HC predictor on learning slope for each level of the other predictor (Clavel, 2015; Fig. 3). The effects of both HC and WM factors on rate of verbal learning is only apparent at higher levels of the other. That is, the positive effect of HC volume on the learning slope factor is only apparent at values of the WM factor above the sample mean. Moreover, the effect of WM on the verbal learning slope factor is only significant in individuals with HC factor scores above the sample mean (Fig. SM3). We confirmed this by re-specifying the model to include additional constraints to test the simple slope of HC on verbal learning separately for WM factor standardized values of ± 0.5 (Clavel, 2015). Model results showed that whereas the low slope of HC on the learning factor at –0.5 on the WM factor was nonsignificant (est. = 0.050, *p* = .841; 95% CI = –0.436 to 0.535), the high slope was significant (est. = 0.536, *p* = .015; CI = 0.105 to 0.967). The positive relationship between HC volume and verbal learning rate was only apparent among those with higher FA in limbic WM.

**Figure 3.**
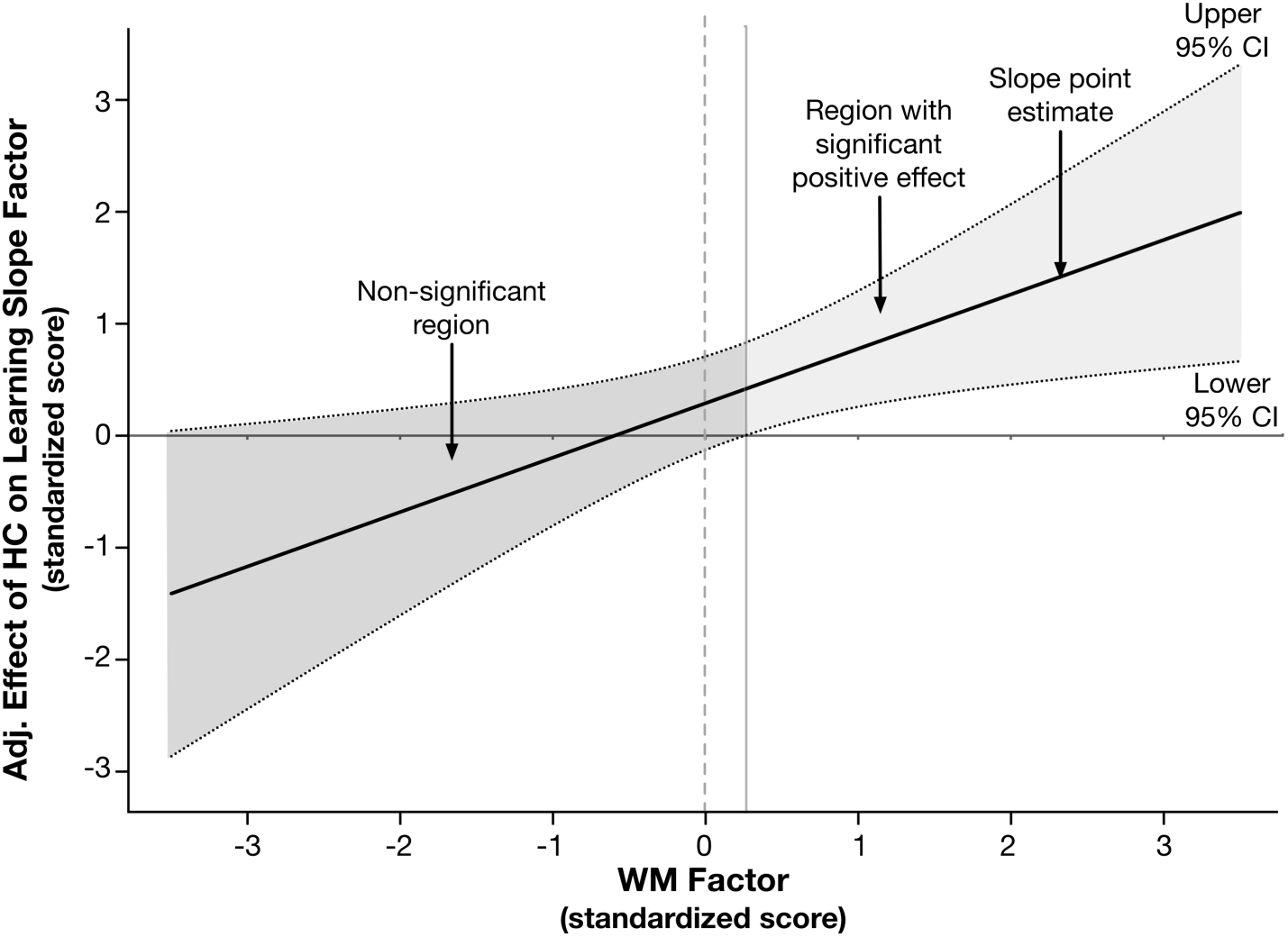
Johnson-Neyman plot illustrating the decomposed interaction to test the moderating effect of limbic white matter FA (WM) on the effect of hippocampal volume (HC) on the verbal learning slope factor. The x-axis represents the continuous moderator – here, the standardized white matter (WM) factor score, and the y-axis represents the effect of the hippocampus (HC) latent factor in the latent interaction on the verbal learning slope parameter, adjusted for other model parameters. The solid regression line reflects the association between the adjusted effect of the hippocampal factor on the learning slope factor, as a function of level in the WM factor. The dotted lines represent the upper and lower 95% confidence band around the regression slope. The solid horizontal line at y = 0, and the dotted vertical line at x = 0 are superimposed to assist with interpretation. Regions where the confidence bands overlap with y = 0 indicates the levels of the x-variable in which the effect represented by the regression slope are not significant; this is denoted by dark gray shading. The confidence bands overlap with zero until the WM factor score is slightly greater than 0.15, demonstrating that the adjusted effect of HC volume on learning is only apparent at non-negative values of the WM factor (i.e., area with lighter gray shading).

To further probe the moderation effect, we saved the standardized factor scores from the model using the factor score regression method. Subsequently, we subdivided the sample distributions for the standardized WM and HC factors into tertiles and used these to examine the results of Johnson-Neyman plots. Our objective for these follow-up analyses was to identify whether different WM×HC patterns in this population-based sample might further qualify differences in learning. Examining bootstrapped (1000 draws) zero-order correlations between learning slope and HC volume by three different levels of WM (Fig. 4) showed individuals in the lowest tertile of FA in limbic WM exhibit negative associations between HC volume and learning slope (*r* = –0.221, *p* = .016; 95% CI = –0.381 to –0.036), which differed from those both in the middle WM factor tertile (*r* = 0.261, *p* = .005; 95% CI *=* 0.084 to 0.428) and in the third WM factor tertile representing highest FA (*r* = 0.392, *p* < .001; 95% CI *=* 0.265 to 0.497).

**Figure 4.**
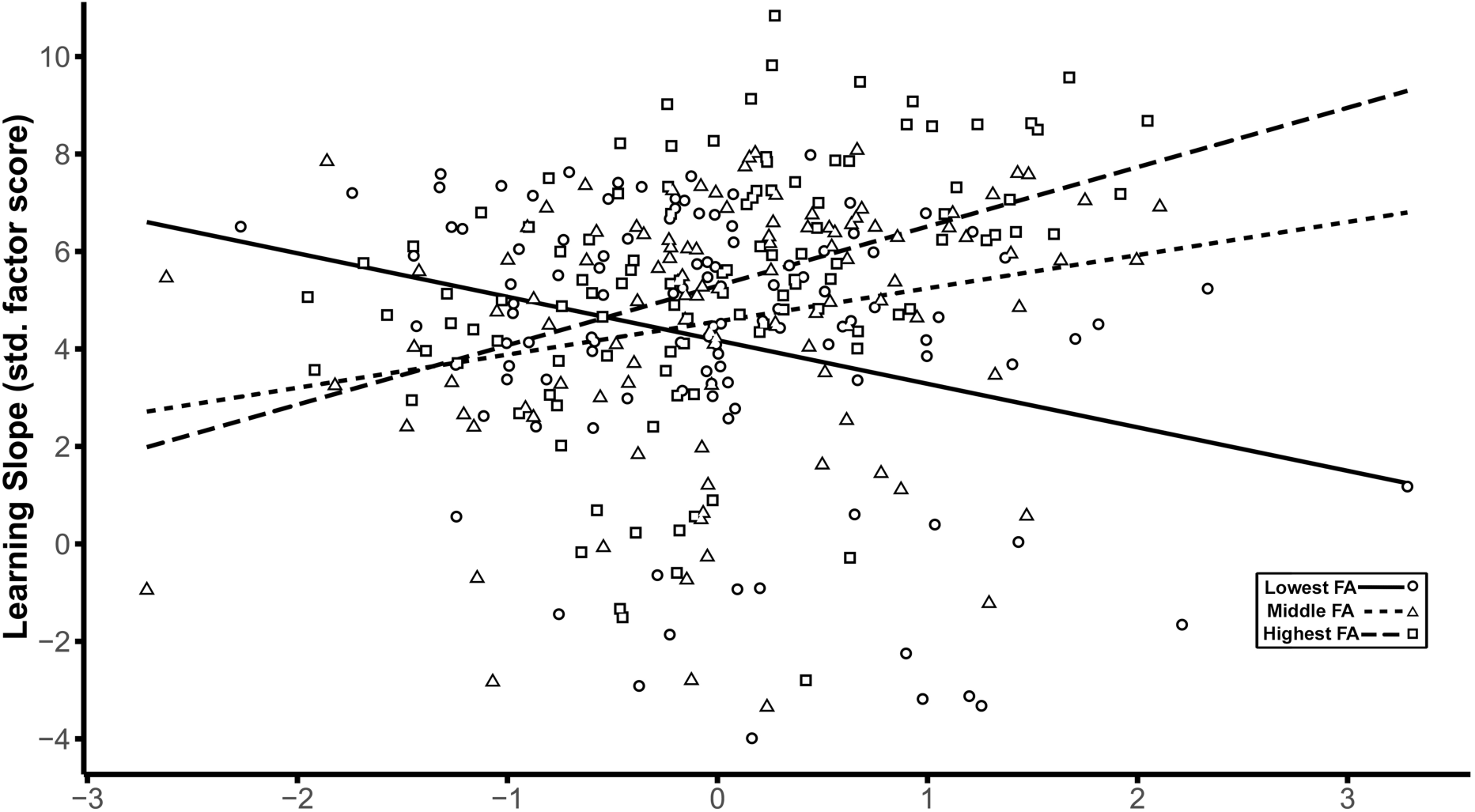
Decomposition of the effects of the latent interaction between hippocampus (HC) and white matter (WM) on the latent factor representing the slope across learning trials based on WM tertiles. The scatter plot shows the HC factor score on the x-axis plotted against the learning slope factor on the y-axis, with linear smoothers fitted separately for each of the three tertiles of the WM distribution. Scales for both axes are depicted using standardized scores. Separate symbols and fitted regression lines represent each of the three tertiles of the WM distribution representing low, middle, and high FA values. Greater HC volume is associated with higher learning slope only in the middle (short-dashed regression line and triangle symbols) and highest tertiles (long-dashed regression line and square symbols) of the WM factor. For the lowest tertile of WM (solid regression line and circle symbols), higher HC volume is associated with lower learning rate.

Next, because the learning slope and intercept factors were not significantly related in the total sample, we inquired whether this association also might jointly depend on HC and WM characteristics. We evaluated differences in the correlation between the slope and intercept factors across the different levels of WM, separately for each tertile of the HC factor distribution. Among participants in the tertile for largest HC, higher learning intercept was associated with less positive slope in those with the highest FA (*r* = –0.248, *p* = .154; 95% CI *= –*0.481 to – 0.020); however, this relationship was positive in both the middle (*r* = 0.522, *p* = .001; 95% CI *=* 0.251 to 0.718), and lower WM tertiles (*r* = 0.261, *p* = .131; 95% CI *= –*0.001 to 0.485). Thus, whereas higher initial recall performance left less room for improvement across learning trials in those with the most robust brain parameters, intercept served as a positive correlate of learning in participants whose brain parameter estimates were near or below the sample mean.

### Covariate models

Although the LMS model without covariates (i.e., the ‘No Covariates’ model in Table 1) provided the primary findings of interest, we repeated the modeling process to assess how these effects are influenced by three relevant demographic covariates: years of age, and educational attainment, both centered at their respective sample means, and participant sex. Initially, we tested the inclusion of covariates in the combined model by specifying paths from each covariate to each of the nine latent factors for HC subfields, WM tracts, and the intercept and slope, which proved an acceptable fit (Table SM2). In addition, covariances between learning slope and WM brain parameters (i.e., FA in fornix and CBH), remained significant in the combined model, with the inclusion of covariates (for both, *p* < .05).

Next, we respecified the second-order models to include the paths from the three covariates to the latent factors for WM, HC, and learning slope. Model fit was acceptable across all second-order covariate models (Table SM5). In the second-order model with all three covariates, the path from the WM factor to learning slope remained significant (estimate = 0.172, *p* = 0.013; 95% CI = 0.055 to 0.765). However, following estimation of the LMS covariate model to test the latent interaction between WM and HC factors on learning, the direct effect from WM to learning rate was no longer significant (estimate = 0.227, *p* = 0.256; 95% CI = –0.165 to 0.620). Of note, however, the path from the HC×WM latent interaction term to the learning rate factor remained significant (estimate = 0.483, *p* = .013; 95*%* CI = 0.102 to 0.864). The LMS covariate model also showed that older age was associated with lower learning slope (estimate = –0.530, *p* = .011; 95*%* CI = –0.937 to –0.124) and with smaller HC (estimate = – 0.267, *p* < .001; 95*%* CI = –0.399 to –0.135). Model results also revealed a significant effect of sex on the learning intercept factor (estimate = 0.592, *p* = .011; 95*%* CI = 0.137 to 1.046) showing superior recall performance by women over men.

Next, we evaluated the differences in levels of covariates across the tertiles of the WM and HC factor distributions. Separately, for each of the three HC factor tertiles we evaluated one-way ANOVAs with age as the dependent variable and WM tertile as the independent variable. The model for the lowest tertile of the HC factor yielded an effect of WM tertile on age, *F*(1,110) = 4.837, *p* = .030, which was rendered nonsignificant following Bonferroni correction for multiple comparisons across the three models. We repeated this process to evaluate differences in educational attainment. The ANOVA revealed a significant effect of WM tertile on education only for the second tertile of HC, *F*(1,110) = 9.564, *p* = .003. Post hoc Student’s t-tests showed that among the participants in the middle tertile of the HC factor, those with lowest WM factor scores had fewer self-reported years of formal education than those with the highest limbic FA: *t*(72) = –3.060, *p* = .003.

## Discussion

We used latent factor modeling to evaluate the relationships between multiple limbic structures and learning in a large, population-based cohort study of older adults. The present study yielded several notable results concerning associations between limbic WM microstructure, HC subfield volumes, and verbal learning. First, a latent factor formed from FA in limbic WM regions and uncinate was consistently associated with faster rate of learning. Moreover, the latent factor representing volume of the hippocampus was not significantly related to learning rate in the total sample. However, evaluating the latent interaction between HC and WM factors revealed an important moderation effect: hippocampal volume was only positively related to learning rate in older adults with more coherent diffusion in limbic WM, possibly reflecting more intact WM microstructure. In contrast, larger hippocampal volume was associated with lower learning rate for individuals with lower WM anisotropy. This has substantial implications for the use of HC volume as a biomarker of brain and cognitive aging.

Van Petten’s (2004) meta-analysis of the relationship between hippocampal volume and memory notes substantial heterogeneity in this association among older adults. The present findings offer one possible explanation for some of this variation. Indeed, Van Petten notes that data from one study of population neuroimaging supported a weak but significant association between total hippocampal volume and immediate and delayed verbal recall (Hackert et al., 2002), with an age-residualized effect of *r* = 0.12. However, that review found that smaller and more selectively sampled study cohorts were often more likely to report the positive association between HC volume and memory in older adults. In comparison, the present population-based cohort study of aging more closely resembles the Rotterdam Study, in which the age-residualized effects of HC volume on memory were rather modest. Thus, population neuroimaging studies that include less selectively screened samples should evaluate the associations between HC volume and memory as conditioned on differences in WM microstructure.

Learning rate has been previously associated with total HC volume in older adults with and without memory impairments (Bonner-Jackson et al., 2015). However, this is the first study to link differences in limbic WM with learning, modeled as a growth function. HC afferent and efferent pathways via the fornix and cingulum play a crucial role in mnemonic encoding and recall (Aggleton and Brown, 1999), underlining the need to examine structural connections beyond the HC (Aggleton, 2014). Prior reports evaluating the combined associations of limbic WM diffusion parameters and HC volumes on episodic memory show mixed effects. Whereas higher FA in CBH and fornix has been linked with better episodic memory, the relationships of total HC volume are inconsistent (Ezzati et al., 2015; Metzler-Baddeley et al., 2011a). We found that higher FA in the ventral (i.e., parahippocampal) portion of the cingulum bundle and the fornix was consistently associated with higher learning rate. Moreover, the association between HC volume and learning rate was positive in individuals with higher FA in limbic WM; however, this relationship was negative in those with low limbic FA.

These results support the notion that the intercept and slope of learning may reflect different demographic factors like age, sex, and education, as well as differences in other cognitive abilities including verbal knowledge, processing speed, and cognitive status (Jones et al., 2005). One possibility is that the different patterns of WM and HC reflect different genetic and life course influences. We found higher educational attainment was associated with more coherent limbic WM microstructure in those in the middle tertile of HC. However, whether this might also serve as an indicator of risk for subsequent decline will require further analysis with longitudinal data. Future studies might also benefit from applying non-parametric approaches to identify non-linear moderation patterns like local SEM (Hildebrandt et al., 2009; Hülür et al., 2011) or SEM trees (Brandmaier et al., 2013). For example, local SEM could be used to move a window over all participants sorted by WM factor values and then plot estimated model parameters over WM factor values.

The present findings also highlight the utility of SEM latent factor approaches for modeling relationships between multiple neural correlates and cognitive measures as latent factors, free from inherent measurement error. This also permits simultaneous estimation of associations between related factors, while precluding the need to correct for multiple comparisons. To the best of our knowledge, this is the first time that the hippocampus has been modeled in this fashion – as a second-order latent factor formed by individual subregional factors. Such an approach may provide a more reliable volumetric estimate of hippocampal structural integrity, particularly in comparison to age-biased estimates of single volumetric indicators from automated segmentation procedures (Wenger et al., 2014).

Furthermore, specifying the latent interaction between the HC and WM latent factors resulted in a better model fit and explained more variance in learning rate. There are a limited number of valid statistical approaches for demonstrating such differential patterns of relationships in cross-sectional data. Mediational approaches are sometimes used to model more complex relationships between brain regions, age and cognition (Foster et al., 2019; Metzler-Baddeley et al., 2019; Salthouse, 2011). Despite violating essential assumptions of temporal ordering necessary to test causal relationships, this nevertheless points to an important modeling need – showing that associations between two variables vary across levels of a third. Moderation approaches are more appropriate for these types of cross-sectional data, and as we show, can illuminate new patterns of brain-cognition relations in the population.

### Limitations and Directions for Future Work

The results of the present study need to be interpreted in light of its limitations. First, the present data are cross-sectional and hence cannot reveal the order and directionality of age-related changes (Lindenberger et al., 2011). Second, we chose to include participants with MMSE scores of 25 and 26, raising the possibility that a small number of participants may have been in the process of developing dementia. There were also several notable technical limitations. First, higher b-values and multi-shell diffusion MRI data can improve resolution of crossing fibers and we recommend their use in future studies. Second, we used aggregate values of WM parameters across tracts of interest, which does not permit more specific anatomical localization of possible effects in cerebral WM. Future studies should try to discern whether specific tract segments are differentially associated with learning and memory (Colby et al., 2012). As the number of brain variables of interest grows (e.g., many regions of interest, or even voxel-level analyses), one may consider statistical approaches that appropriately deal with situations with large number of predictors and relatively small sample sizes, such as regularization (Jacobucci et al., 2019). Also, the 2mm slice thickness associated with the high-resolution structural imaging sequence for HC subfield volumetry used in this study may have come with cost of inducing greater partial volume artifacts. In addition, HC subfield measurement was limited to the body. Although some published methods permit segmentation of the head and tail, this may simply introduce further methodological heterogeneity (Yushkevich et al., 2015b). Work currently in progress should help extend valid segmentation of HC subfields to head and tail of the HC using a harmonized protocol (Wisse et al., 2017).

Last, there are also assumptions and limitations associated with specifying interactions in latent space (Moosbrugger et al., 1997). One concern is that established estimation methods impose potentially problematic assumptions regarding orthogonality of error structures (Little et al., 2006). However, most published work with such SEM approaches for testing latent factor moderation specify exogenous latent factors based on unreliable observed variables. Here, we used a second-order factor, and although this may be a viable method for circumventing such concerns, such an approach has not been compared before with other latent moderation approaches. Thus, further work is needed to establish better practices for estimating interactions between continuous factors. Moreover, future studies should also compare changes in WM and HC measures as correlates of longitudinal changes in learning (Bender and Raz 2015; Bender et al., 2016). It is unclear why the paths from both WM and HC factors to the learning slope were attenuated following inclusion of the age covariate, but that their interaction was not. Further work is needed to investigate the possibly differentially age-related mechanisms that underlie HC and WM and their interaction.

## Conclusion

In the present study, we delineated multimodal neural correlates of verbal learning in older adults, including specific limbic WM fiber tracts and HC subregions. We show hippocampal volumetric associations with verbal learning are dependent on the levels of FA in limbic WM fiber tracts. Given that the present sample was unimpaired and did not widely differ in age, we consider this result as encouraging (cf. Salthouse, 2011), while recognizing that it needs to be replicated and extended in future cross-sectional and longitudinal investigations.

These findings also suggest future studies should account for differences in WM microstructure when considering total hippocampal volume as a correlate of learning and memory in older adults.

## Acknowledgments

This study was supported by the Strategic Innovation fund of the Max Planck Society to UL (67– 11HIPPOC), and by the National Institutes of Health grants (P41EB015902, R01AG042512) to OP. SK was supported by grants from the German Science Foundation (DFG KU 3322/1-1, SFB 936/C7). This project is also part of the BMBF funded EnergI consortium (01GQ1421B) We thank Michael Krause for technical assistance in this study, Dhaval Gandhi for assistance with manuscript preparation, and Ahnalee Brinks for suggestions regarding data visualization.

The authors have no conflicts of interest to declare.

## Supplementary Materials

### Methods

#### ROI creation

Three regions were used to specify inclusion of fornices: a seed mask was placed in the coronal plane on the body of the fornix and transverse-oriented inclusion masks were drawn at the level of mammillary bodies and, and bilateral masks were drawn on the coronal plane at crus of the fornix, one just posterior hippocampal body and one more superior. Exclusion masks were drawn 1) on the coronal plane, two slices anterior to the fornix, 2) on the coronal plane, in the splenium of the corpus callous (CC), and 3) anterior to the columns of the fornix, and 4) inferior to the mammillary bodies. The CBD was defined by a single seed ROI placed in the middle of the dorsal cingulum, with regions of inclusion and termination anterior to the genu of the CC and posterior to the CC splenium. Exclusion masks were liberally placed dorsal, rostral, and caudal to the CBD to limit the tract to its core projections. A seed ROI for CBH was placed in the coronal plane, in middle of the tract as visualized in the parahippocampal gyrus with regions of inclusion and termination at drawn on the coronal plane at the uncal apex in the hippocampal head (anterior) and posterior termination at the inferior aspect of the splenium of the corpus callosum, drawn on the transverse-oriented image. Uncinate fasciculus was determined using a region of inclusion at the external capsule on the coronal plane on the slice in which the temporal lobe and frontal lobe become contiguous with regions of inclusion at the anterior temporal lobe and in the prefrontal lobe. Regions of exclusion were liberally applied to mask out additional streamlines. Fornix tracts were subsequently edited to restrict streamlines to the posterior aspect, covering the crus and posterior body.

#### Spatial transformation of masks to native space

The spatial transformation matrices produced by DTI-TK during registration to template space were inverted and combined. We sampled the s-form matrix from the original diffusion volume prior to nonlinear registration and used this information to reorient the deprojected masks from neurological to radiological orientation, and back to the native coordinate space.

#### Inspection of streamlines

In 18 cases, the rater adjusted the inclusion masks for CBD and CBH, or swapped seed and inclusion masks. Cases with fewer than 15 streamlines were deemed unreliable and treated as missing values. Streamlines for fornix were particularly vulnerable to producing too few streamlines, and cases with less than 15 streamlines in left, right, or bilateral fornix were produced in approximately 10% of the sample. The numbers of cases assigned as missing values, varied by region and hemisphere, and the final totals of cases with missing streamlines were as follows: CBD-left = 8, CBD-right = 8, CBH-left = 16, CBH-right = 23, fornix-left = 31, fornix -right = 44, UF-left = 8, UF-right = 8.

#### Hippocampal subfield segmentation

We used the Automated Segmentations of Hippocampal Subfields (ASHS; Yushkevich et al., 2015; Yushkevich et al., 2010) software with a customized atlas for HC subfield morphometry (Bender et al., 2018). The customized atlas was built using a modified version of the manual demarcation and tracing rules described previously (Bender et al., 2013; Daugherty et al., 2016), and includes a slightly more lateral placement of the SUB-CA1/2 boundary as a compromise of that boundary placement in different atlases included in commonly used HC subfield segmentation software (Iglesias et al., 2015; Yushkevich et al., 2015). This atlas was built from a lifespan sample, and included data from 10 children and adolescents (age range = 7–13 years; mean age = 10.08, SD = 2.64 years; 50% female), four young adults (age range = 22–24 years; mean age = 23.00, SD = 0.82 years; 50% female), and 14 older adults (age range = 62–78 years; mean age = 69.64, SD = 4.63 years; 50% female). To ensure ASHS demarcation was performed on the full extent of the HC body, we used an extended atlas (Bender et al., 2018). The extended atlas was built from manually demarcated data in which the ranges of inclusion were extended beyond anatomical landmarks normally designated for manual segmentation procedures. Thus, images included for atlas building by tracing subfields on one to two additional anterior slices and one to two additional posterior slices to the manually defined ranges, depending on visibility of subfields. In anterior slices, any visible tissue from the long digitation of the HC head was not included and demarcation was limited to clearly apparent HC ‘body-like’ regions on slices anterior to the uncal apex. Automated segmentation failed or produced errors in 36 out of 337 cases (10.68%) included for analysis, and these were treated as missing values in subsequent analyses.

#### Manual range determination

We separately determined the ranges of slices for inclusion in each HC subfield region of interest (ROI) for left and right hemisphere. The first slice following the uncal apex, and on which the long digitation of HC head was no longer visible and did not exhibit partial volume artifacts served as the anterior limit of HC body. The *penultimate* slice on which the lamina quadrigemina (LQ) was visible served as the posterior limit for inclusion. We allowed for hemispheric differences in posterior range if only left or right LQ was visible on the final slice, including presence of a partial volume effect. Using a custom Bourne shell script, we truncated the output from ASHS to the individualized, manually-determined ranges.

#### ICV correction

As described previously (see Bender et al., 2013 for a complete description) Standard-space masking was applied to remove non-brain tissue, and a fractional intensity threshold of 0.2 and we used the -A option for the ‘betsurf’ feature for estimation of skull surfaces (Jenkinson et al., 2005). An experienced operator (ARB) reviewed the results and identified 11 cases in which the procedure produced holes in the ICV brain mask, and filled the holes using tools in ITK-SNAP. We used ICV values sampled from the outer skull mask to adjust HC subfield volumes for head size.

### Data Analysis

#### Latent growth model (LGM)

We specified a model with latent factors for intercept and slope. For all trials, the factor loadings for the intercept were all fixed at 1. The factor loadings for the 5 trials were estimated using a latent basis free model approach (McArdle, 1986) in which the first and last indicator factor loadings are fixed at 0 and 1, respectively, and the remaining factor loadings are freely estimated.

### Brain Parameter Confirmatory Factor Analyses (CFA)

#### Brain CFAs

All CFA models for HC subfield volumes converged with acceptable fit according to most indices. Initial estimation of the three–factor model showed negative residual variance for the left CA3/DG indicator. We addressed this by fixing the residual variance for this indicator to zero for all subsequent modeling steps, which resulted in unstandardized factor loading of 1 for this indicator. As shown in Table SM1, the standardized factor loadings for the remaining HC subfield factor indicators were estimated between 0.711 (right CA3/DG) and 0.962 (left CA1/2). The comparison of model fit indices for one- and four–factor models showed the four–factor model to provide the best fit (Table SM2). Model fit in the four–factor model was improved by specifying covariances between indicators for subfields within each hemisphere. The four–factor model showed significant associations between all HC subfield factors, ranging from moderate to strong.

Initially, for both HC subfield volumes and DTI tracts, we compared the model fit of two different approaches: (1) separate latent factors per region or tract with the correlations between latent factors freely estimated, and (2) all indicators loading onto a single factor (SM Fig. 2 similar to a first principle component). For both HC subfield volumes and each WM measure, we assessed which measurement model (individual latent factor or single factor) best fit the data.

Comparison of model fit for the WM factor models showed poor fit by the one-factor model, and excellent fit by each of the four–factor models (see Table SM2).

## Results

### Intra-domain brain associations

Consistent with our expectations that factors within the respective domains (i.e., WM or HC subfield) would be highly correlated, we observed significant associations between all HC subfield factors, and between all WM factors for FA, (Table SM2).

---

**Table SM1.** Standardized factor loadings from single-factor latent models

|  | Left | Right |
| --- | --- | --- |
| *HC Subfield* |  |  |
| SUB | 0.918 | 0.698 |
| CA1/2 | 0.962 | 0.714 |
| CA3/DG | 0.998 | 0.711 |
| *FA* |  |  |
| CBD | 0.902 | 0.825 |
| CBH | 0.785 | 0.920 |
| Fornix | 0.783 | 0.416 |
| UF | 0.827 | 0.776 |

Notes: SUB – subiculum; CA1/2 – aggregated ROI for CA1 and CA2 subfields; CA3/DG – aggregated ROI for CA3 and DG; CBD – dorsal cingulum bundle; CBH – hippocampal cingulum bundle; UF – uncinate fasciculus; FA – fractional anisotropy

**Table SM2.** Goodness of Fit Statistics for CFAs and Covariance Models

| Model | $\chi^2(df)$ | $p$ | $\chi^2/df$ | CFI | RMSEA | RMSEA 90% CI | SRMR |
| --- | --- | --- | --- | --- | --- | --- | --- |
| *Verbal Learning Memory Test (VLMT) LGM* |  |  |  |  |  |  |  |
| Linear slope | 182.129 (10) | .000 | 18.213 | 0.839 | 0.226 | 0.039–0.120 | 0.130 |
| Latent basis | 42.554 (7) | .000 | 6.079 | 0.967 | 0.123 | 0.089–0.159 | 0.074 |
| *HC Subfield Volume* |  |  |  |  |  |  |  |
| 1-Factor | 279.547 (6) | .000 | 46.591 | 0.791 | 0.389 | 0.351–0.429 | 0.064 |
| 3-Factor | 3.563 (4) | .468 | 0.891 | 1.000 | 0.000 | 0.000–0.083 | 0.020 |
| *DTI - FA* |  |  |  |  |  |  |  |
| 1-Factor | 288.511 (20) | .000 | 14.426 | 0.704 | 0.202 | 0.181–0.223 | 0.086 |
| 4-Factor | 9.437 (14) | .802 | 0.674 | 1.000 | 0.000 | 0.000–0.035 | 0.013 |
| *Combined Model* |  |  |  |  |  |  |  |
| HC-WM | 156.026 (136) | .115 | 1.147 | 0.994 | 0.021 | 0.000–0.035 | 0.030 |
| HC-WM-Covs‡ | 199.518 (150) | .004 | 1.330 | 0.985 | 0.031 | 0.018–0.042 | 0.036 |
| *Second-order factor model* |  |  |  |  |  |  |  |
| Covariances only | 171.26 (119) | .001 | 1.439 | 0.984 | 0.036 | 0.023–0.048 | 0.043 |

Notes: ‡: Covariate model including direct paths to each latent factor from observed covariates: participant years of age and educational attainment (centered at sample mean) and sex.

**Table SM3.** Comparison of factor loadings in latent growth models

| Model | Factor Loadings by VLMT Trial |  |  |  |  | Model Fit |  |  |
| --- | --- | --- | --- | --- | --- | --- | --- | --- |
|  | 1 | 2 | 3 | 4 | 5 | RMSEA | CFI | SRMR |
| Latent Basis Model* | 0.000 | 0.452 | 0.698 | 0.889 | 1.000 | 0.118 | 0.961 | 0.097 |
| Linear Slope | 0.000 | 1.000 | 2.000 | 3.000 | 4.000 | 0.238 | 0.839 | 0.139 |

Notes: Latent basis model = Factor loadings for trial 1 fixed to 0 and for trial 5 fixed to 1, and all other loadings are freely estimated. Note that the difference in the last factor loading between models merely reflects a rescaling of the latent slope variable but does not affect model fit. \* indicates factor loadings used in subsequent modeling steps.

**Table SM4.** Significant associations between parameters within brain domain

|  | Association | Estimate | S.E. | *p* | 95% CI |
| --- | --- | --- | --- | --- | --- |
| HC Subfields |  |  |  |  |  |
|  | CA1/2↔SUB | 0.649 | 0.042 | .000 | 0.560–0.734 |
|  | CA3/DG↔SUB | 0.629 | 0.040 | .000 | 0.548–0.705 |
|  | CA3/DG↔CA1/2 | 0.820 | 0.024 | .000 | 0.771–0.864 |
| WM fractional anisotropy (FA) |  |  |  |  |  |
|  | CBH↔CBD | 0.554 | 0.050 | .000 | 0.460–0.639 |
|  | Fornix↔CBD | 0.223 | 0.080 | .005 | 0.059–0.408 |
|  | Fornix↔CBH | 0.365 | 0.090 | .000 | 0.166–0.578 |
|  | UF↔CBD | 0.552 | 0.051 | .000 | 0.450–0.645 |
|  | UF↔CBH | 0.585 | 0.051 | .000 | 0.445–0.690 |
|  | UF↔Fornix | 0.303 | 0.087 | .000 | 0.100–0.513 |

Notes. SUB = subiculum, CBD = dorsal cingulum bundle, CBH = hippocampal cingulum bundle, UF = uncinate fasciculus.

**Table SM5.** Fit indices for 2^nd^ order models without latent interactions

| Model | $\chi^2$ (df) | $\chi^2 / df$ | *p*-value | CFI | RMSEA | SRMR |
| --- | --- | --- | --- | --- | --- | --- |
| No Covariates | 221.776 (144) | 1.54 | .000 | 0.977 | 0.040 | 0.051 |
| Age | 250.185 (161) | 1.55 | .000 | 0.974 | 0.041 | 0.051 |
| Age, Sex | 299.782 (176) | 1.70 | .000 | 0.964 | 0.046 | 0.053 |
| Age, Sex, Edu | 309.887 (191) | 1.62 | .000 | 0.959 | 0.042 | 0.052 |

Notes: Results shown for reduced model, specified without latent intercept factor as a dependent variable.

**Table SM6.**

| Model | *LL-Orig* | *LL-LI* | *LLR* |
| --- | --- | --- | --- |
| No Covariates | -530.08 | -527.19 | 5.776* |
| Age | -502.81 | -500.04 | 5.524* |
| Age, Sex | -501.78 | -499.15 | 5.258* |
| Age, Sex, Edu | -1241.20 | -1238.50 | 5.404* |

Notes: LL: log-likelihood value. Orig: Original model without latent interaction. LI: Latent moderated structural equation model specifying the interaction between second order factors for hippocampal volume and limbic white matter fractional anisotropy. LLR: Log-likelihood ratio test, a two-tailed test of differences in fit. LLR statistic shows significant differences between models including latent interaction and those that do not, with lower log-likelihood indicating better fit.

**Figure SM1.**
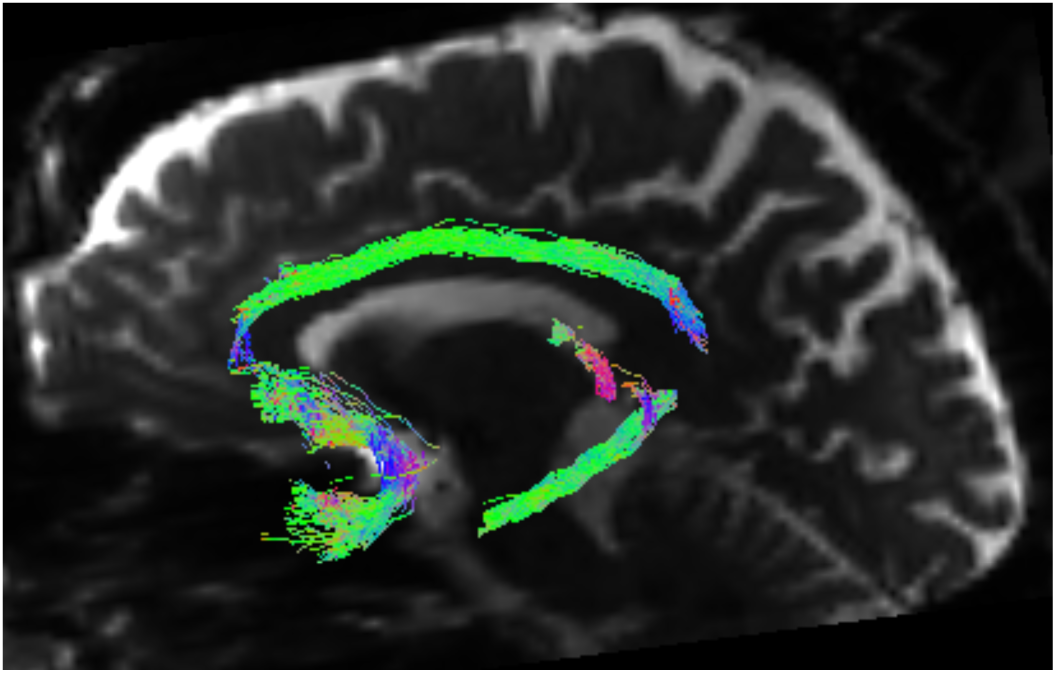
Illustration of constrained spherical deconvolution diffusion tractography using MRtrix3. Using deterministic fiber tractography we sampled free water-corrected fractional anisotropy from streamlines representing canonical limbic system white matter fiber tracts: uncinate fasciculus, dorsal and parahippocampal cingulum bundle, and posterior fornix.

**Figure SM2.**
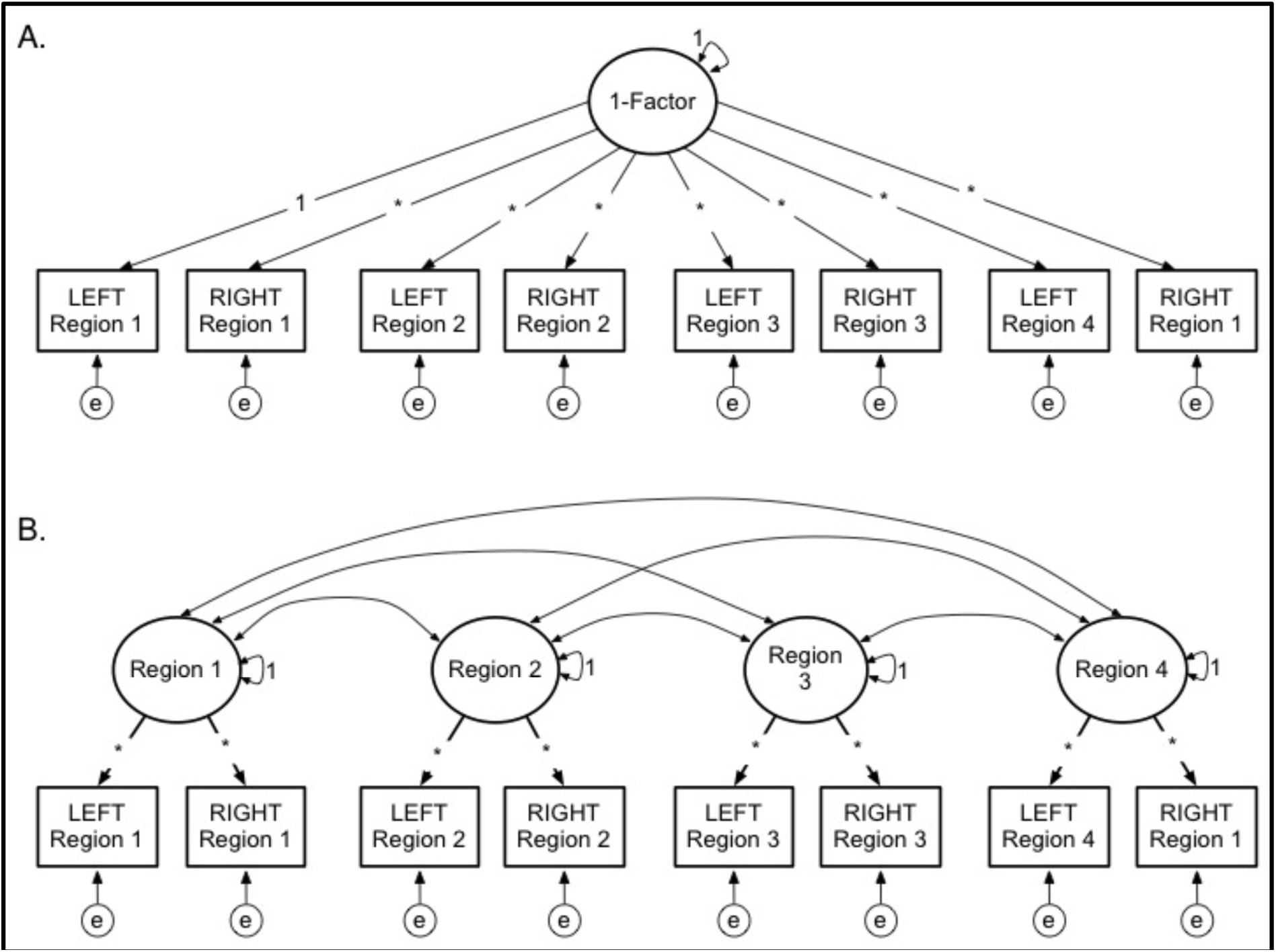
Alternative measurement models for DTI parameters and hippocampal subfields. A. One-factor model with all indicators loading onto a single factor. B. Four-factor model, with dual indicators representing left and right hemispheres for each latent factor representing individual, bilateral anatomical regions (i.e., WM tract or HC subfield volume).

**Figure SM3.**
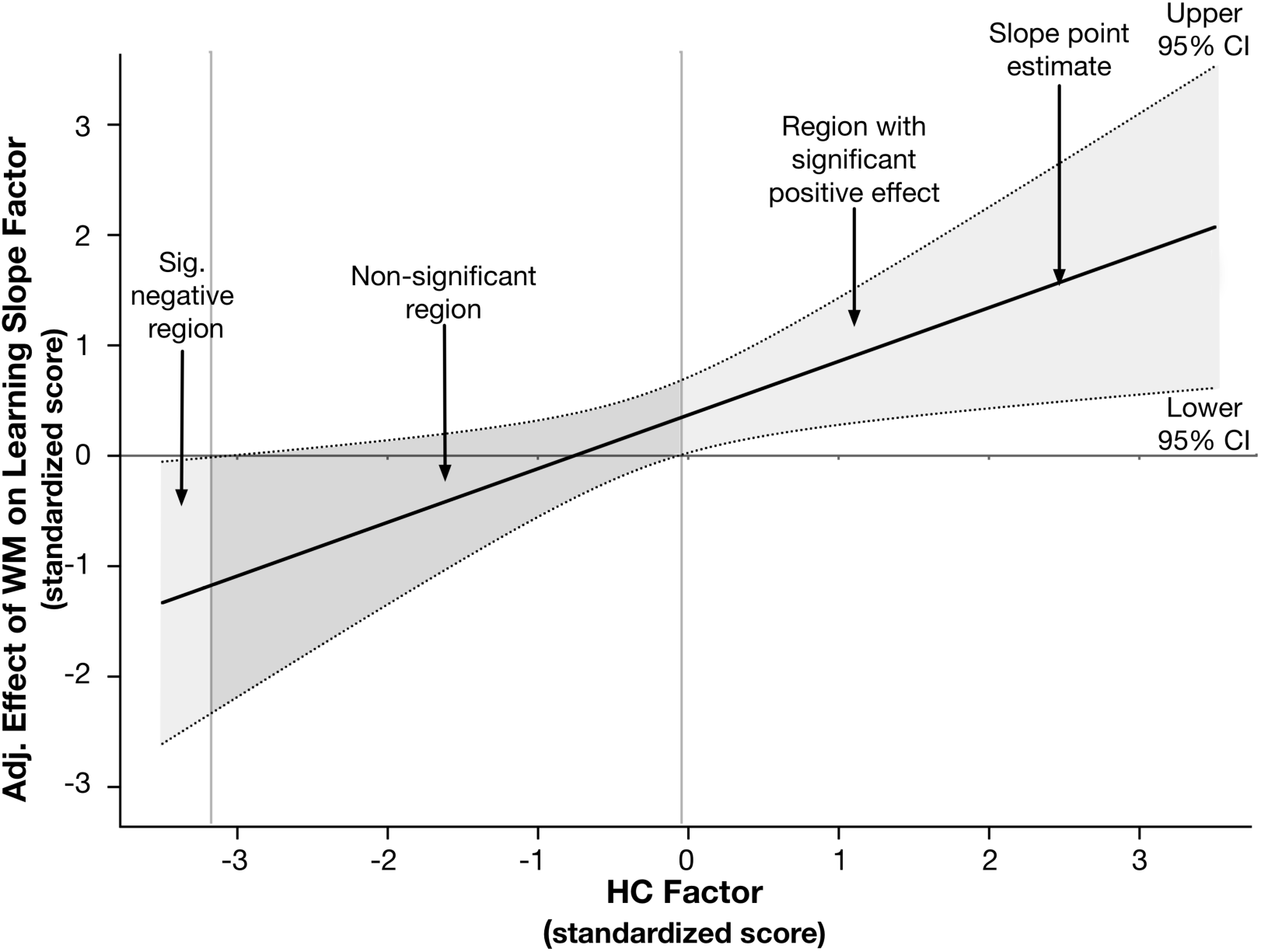
Johnson-Neyman plot illustrating the decomposed interaction to test the moderating effect of hippocampal volume (HC) on the effect of limbic white matter (WM) fractional anisotropy on the verbal learning slope latent factor. The x-axis represents the continuous moderator – here, the standardized HC factor score, and the y-axis represents the effect of the WM latent factor in the latent interaction on the verbal learning slope parameter, adjusted for other model parameters. The solid regression line reflects the association between the adjusted effect of the WM factor on the learning slope latent factor, as a function of level in the HC factor. The dotted lines represent the upper and lower 95% confidence band around the regression slope. The solid horizontal line at y=0, and the dotted vertical line at x=0 are superimposed to assist with interpretation. Regions where the confidence bands overlap with y=0 indicates the levels of the x-variable in which the effect represented by the regression slope are not significant. The confidence bands overlap with zero until the HC factor score is slightly greater than 0, demonstrating that the adjusted effect of WM volume on learning is only apparent at non-negative values of the HC factor.

## Notes

#### Summary of Updates

This updated includes minor changes to post hoc models decomposing the effect of the interaction between second-order factors representing hippocampal volume and white matter fractional anisotropy on the slope of verbal learning, as well as changes to the figure supporting that analysis.

